# Graph-Based Method for Anomaly Detection in Functional Brain Network using Variational Autoencoder

**DOI:** 10.1101/616367

**Authors:** Jalal Mirakhorli, Mojgan Mirakhorli

## Abstract

Functional neuroimaging techniques using resting-state functional MRI (rs-fMRI) have accelerated progress in brain disorders and dysfunction studies. Since, there are the slight differences between healthy and disorder brains, investigation in the complex topology of human brain functional networks is difficult and complicated task with the growth of evaluation criteria. Recently, graph theory and deep learning applications have spread widely to understanding human cognitive functions that are linked to gene expression and related distributed spatial patterns. Irregular graph analysis has been widely applied in many brain recognition domains, these applications might involve both node-centric and graph-centric tasks. In this paper, we discuss about individual Variational Autoencoder and Graph Convolutional Network (GCN) for the region of interest identification areas of brain which do not have normal connection when apply certain tasks. Here, we identified a framework of Graph Auto-Encoder (GAE) with hyper sphere distributer for functional data analysis in brain imaging studies that is underlying non-Euclidean structure, in learning of strong rigid graphs among large scale data. In addition, we distinguish the possible mode correlations in abnormal brain connections.

## Introduction

The human brain has a complex connection of various parts which dynamically shift during its operation. Therefore, the model and cost of each part can change according to type of its operation in carried out or rest state. the fMRI data exhibits non-stationary properties in the context of task-based studies [1, 2]. Therefore, the analysis of these sections are outmost value and is able to predict the connection factors for each independent profile. Here in, we present a theoretical model based on VAE and graph theory to learn probability distributed of graph that can to extract the data model of tasks from brain regions with a semi-unknown prior knowledge method. We used each tasks-base functional connectivity matrix that were collected in an rs-fMRI experiment, using rs-fMRI data from Alzheimer’s Disease Neuroimaging Initiative (ADNI). Functional connectome analysis is able to reveal biomarkers of individual psychological or clinical traits and describes the pairwise statistical dependencies which exist between brain regions [3]. In this article, we present brain as a graph using functional connectome structures, which allow us to probing and inference about how dynamic changes progress of improvement degree of brain disorder or predict the disease as well as the term brain abnormalities. This paper propose to introduce a framework for feature extraction of the brain graphs which provide across many subjects, for prediction of ambiguous parts of brain. In this method a VAE is developed to make the graph and experiment a Bayesian Von Mises–Fisher (VMF) [4] mixture model as a latent distribution that can place mass on the surface of the unit hypersphere [5] and stable the VAE. Our experiments demonstrate that this method significantly outperform other methods and is a large step forward to inference brain structure. It is capable to handle both homogeneous and heterogeneous graph. Recent studies have shown, geometric deep learning methods have been successfully applied to data residing on graphs and manifolds in terms of various tasks[6,7], for example brain function prediction and its graph expression analysis address the multifaceted challenges arising in diagnosis of brain diseases. Here in, we present a novel method using a graph model in revealing of the relationship between the parts of brain and recover missing parts or no properly function parts.

## Related works

As our approach focuses on completing graph and prediction defective parts of graph with obtained feature of network embedding, we consider some of related fields. In additional, we used a combination of graph convolution VAE to address both recovery and learning problems which can be performed in spectral [8, 9] or spatial domain [10]. D. Xu and et al in [11] construct a graph from a set of object proposals, provide initial embedding to each node and edge while used message passing to obtain a consistent prediction. Simonovsky and Komodakis in [12] used a generative model to produces a probabilistic graph from a single opaque vector without specifying the number of nodes or the structure explicitly. Pan and et al in [13] proposed an adversarial training scheme to regularize and enforce the latent code to match a prior distribution with a graph convolutional Autoencoder. Makhzani in [14] showed an adversarial Autoencoder to learn the latent embedding by merging the adversarial mechanism into Autoencoder for general data but Dai and et al [15] applied the adversarial procedure for the graph embedding. Also in [12] used an encoder with edge condition convolution (ECC) [16] and condition both encoder and decoder which associated with each of the input graph, this method is useful only for generation small graphs.

## Approach

In spite of individual alteration, human brains performed common patterns among different subjects. Therefore, algorithms base on graph are essential tool to capture and model complicated relationship between functional connectivity.in this work, we used a model of graph embedding to convert graph data into a low dimensional and compaction continuous feature space that is able to detect abnormal parts of input graphs [17] which is involved with graph matching and partial graph completion problems. To develop this algorithms need to present a generative model that construct from a Graph Varational Autoencoder with hypersphere distribution [18,19,20]. Partial abnormality can be appear by features train in latent space, considering both first-order proximity, the local pairwise proximity between the vertices in the network, and second-order proximity. This refers to vertices sharing many connections to other vertices that are similar to each other. The work flow of the algorithm, in more details, is showed in figure 1.

**Fig 1.**
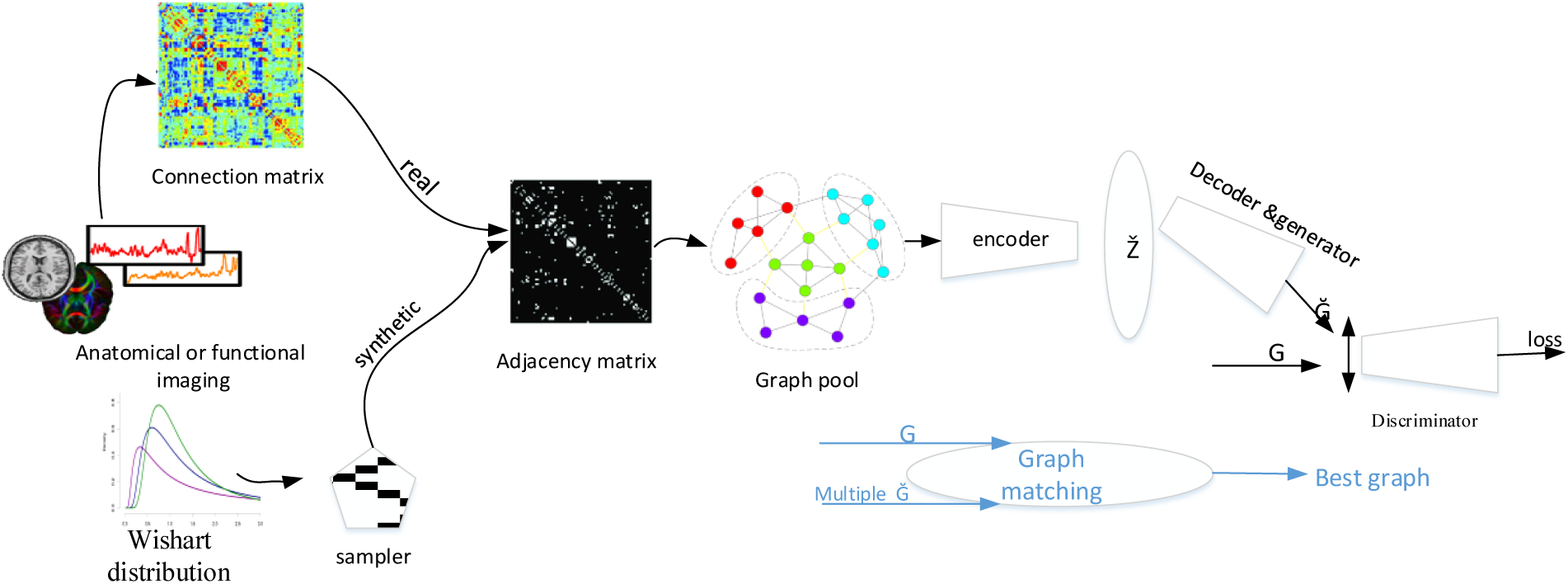
The data flow of the proposed network architecture

### Brain network as a graph

As shown in figure 1, using rs-fMRI data of subjects acquired by preprocessing ADNI dataset to provide an adjacency matrix that encodes similarities between nodes and a feature matrix representing a node’s connectivity profile, to define the input data as an undirected, connected graph G = (V, E, W), which consists of a finite set of vertices V with |V| = n, a set of edges E, and a weighted adjacency matrix W. If there is an edge e = (i, j) connecting vertices i and j, the entry W_ij_ or a_ij_ represents the weight of the edge a_ij>0_, otherwise a_ij_ = 0. For each of n subjects make a data matrix X_n_ ε R ^dn*dy^, where d_y_ is the dimensionally of the node’s feature vector. This structure of fMRI data will be merge to the graph defined. We will show that graph base algorithm on brain connectivity is useful to analyze brain information processing.

### Graph Convolutional Neural Network (G-CNN)

for apply convolution-like operators over irregular local supports, as graphs where nodes can have a varying number of neighbors which can be used as layers in deep networks, for node classification or recommendation, link prediction and etc. in this process we involved with three challenges, a) defining translation structure on graphs to allow parameters sharing, b) designing compactly supported filters on graphs, c) aggregating multi-scale information, the proposed strategies broadly fall into two domains, there is one spatial operation directly perform the convolution by aggregating the neighbor nodes’ information in a certain batch of the graph, where weights can be easily shared across different structures[21,22] and other one is spectral operation relies on the Eigen-decomposition of the Laplacian matrix that apply in whole graph at the same time [23,24,25,26], spectral-based decomposition is often unstable making the generalization across different graphs difficult[10], that cannot preserve both the local and global network structures also require large memory and computation. On the other hand, local filtering approaches [27] rely on possibly suboptimal hard-coded local pseudo-coordinates over graph to define filters. The third approach rely on point-cloud representation [28] that cannot leverage surface information encoded in meshes or need ad-hoc transformation of mesh data to map it to the unit sphere [29].overall, spectral approach has the limitation of graph structure being same for all samples i.e. homogeneous structure, this is a hard constraint, as most of sample graphs in the learning phase has same structures and size for different structures i.e. heterogeneous structures. Then here, we apply the spatial approach that is not obligatory homogeneous graph structure, in turn requires preprocessing of graph to enable learning on it. Therefore we used a method that propose a graph embed pooling. In [30] Graph convolution transforms only the vertex values whereas graph pooling transforms both the vertex values and adjacency matrix. Convolution of vertices V with filter H only require matrix multiplication of the form, υ_out_=Hυ_in_ where υ_in_, υ_out_ ε R^N*N^. the filter H is defined as the k-th degree polynomial of the graph adjacency matrix A;

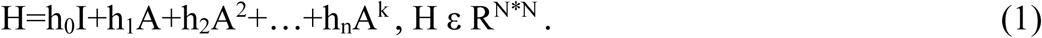

We used the first two taps of H for any given filter.

### Graph Autoencoder (GAE)

GAE is inherently an unsupervised generative model, our model is closely base on framework of Varational Autoencoder by [20,31]. In follow we briefly describe GAE and introduce our method with objectives. For learning both encoder, decoder in the figure 1 to map between the space of graph and their continuous embedding Z ε R^C^, stochastic graph encoder q_Φ_(Z|G) embed the graph into continuous representation Z. given a point in the latent space Z, the graph decoder p_θ_ (G|Z) outputs a probabilistic fully-connected graph Ğ on predefined nodes, where Φ,θ are learned parameters. Reconstruction ability of GAE is facilitated by approximate graph matching for aligning G with Ğ, as well as a prior distribution P(Z) imposed on the latent code representation as a regularization and train GAE via optimization of the marginal likelihood, P(G)_=_ ∫ *Pθ(Z)P*(*G*|*Z*) *dz*, then the marginal log likelihood can be written;

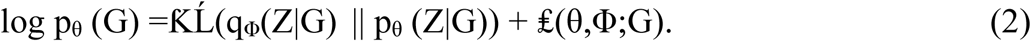

Where Kullback–Leibler (ƘĹ) and are a divergence term in loss function that encourages the Varational posterior and a Varational approximation to the posterior p (Z|G), respectively. Here, we used a hyperspherical latent structure for parameterization of both prior posterior, because one of important limitation using Gaussian mixture is that ƘĹ term may encourage the posterior distribution of the latent variable to collapse in prior or tends to pull the model toward the prior, during approximation the prior, whereas in the VMF case is not such pressure toward a single distribution convergence. Therefore a VMF [32,33] distribution is more suitable for capturing data[20], VMF distribution defines a probability density over points on a unit-sphere also The consequences of ignoring the underlying spherical manifold are rarely analyzed in parts due to computational challenges imposed by directional statistics.

### Geometric deep learning

for graph generation, we employed the GAE to graph G ε R^n*m^ under an unsupervised learning method, our goal is to learn an implicit generative mode that can predict abnormal sections in the graph, of course here we are not sure that close links have similar features to detect unseen deformable and hidden angle of graphs. Our method inspired by [34, 35], and combination from the GAE and generative adversarial network (GAN) that decoder of GAE and generator of GAN have been supportive role. Following the above mentioned items, we used the uniform distribution VMF(0,Ҡ =0) for our prior and approximate p_θ_ (Z | Ğ) with variational posterior q_Φ_(Z|G) = VMF(Z; μ, Ҡ), where μ is mean parameter and Ҡ is a constant, the variational distribution is associated with a prior distribution over the latent variables, our GAE loss combines the graph reconstruction Ĺ_r_=‖ Ğ – G ‖_2_ encouraging concatenation both the encoder-decoder to be a nearly identity transformation, a regularization prior loss measured by the ƘĹ divergence, Ĺ_p_=D _ƘĹ_(q(z|G)‖ P(Z)) and a cross entropy loss Ĺ_2D-GAN_ for GAN, Ĺ _G-GAN_=log D(G)+ log (1-D(G(z))), where D is discriminator as a confidence D(G) of the whether a input graph G is real or synthetic[24]. The total GAE+GAN loss is computed as Ĺ= Ĺ_r_ +λ_1_Ĺ_p_+λ_2_Ĺ_G-GAN_ where λ_1_ and λ_2_ are weights of ƘĹ divergence loss and reconstruction loss. As discussed in above, our desire to focus on graph completion for deformable object classes in brain connectome therefore we used dynamic weight of filtering in each convolutional layer.

### Partial graph completion

once our model GAE-GAN has been trained, the encoder the element of GAN are discarded away, so that the role of the decoder is only as a graph generator that a probabilistic latent space z act as a base for finding the target graphs the same graph prior. At inference, for each space of the latent vector z may represent a few complete graph correspondence a latent vector, then partial graph or deformation graph in the input of system makes a few complete graph in the output, the higher deformation rate in input, the more of graph is generated. Each partial graph represent with the partial adjacency matrix δ that apply it to any graph Ğ generated by our model and explore similarity between their, for finding best compatibility or a latent vector z* which can minimize differences between input and output graph, to provide more geometric insight on the problem. Process to measure similarities among elements of graphs with probing combination of dependences similar unary, pairwise or high-order [36, 37, 38] as well as there are Potentials between reference graph and their counterparts similar to [39,40] That follow a function is used for finding dissimilarity or deformation with a convex optimization problem over the set of doubly stochastic matrices.

### Graph recovery plan

As mentioned above, our goal is the optimal choice a latent vector z* so that minimize dissimilarity of between the partial graph related to a disease brain, G, and the generated graph, Ğ =dec(z);min (Ğ, ζG)

Where ζ denote a rigid transformation, this procedure is performed over z, non-rigid deformation and ζ crosswise. Similar to [39] minimizing the following function is our goal as an objective function;

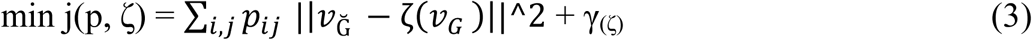

Where γ is a regularization term of geometric transformation ζ: Ğ→G, p is a map for measuring the difference of graph attributes in similarity transformation domain, in each step of optimization a weight matrix measure the degree of deformation on the radial basis function method [39], graph recovery is ill-posed problem that has multiple plausible solution while in this paper we limit the prediction space to only several structures of the graphs.

## Materials and Discussion

In the present study, we employed an rs-FMRI data to construct graphs via adjacency matrix and feed the graphs into our model to exploring functional brain network alteration in patients with Alzheimer’s disease (AD).GAE was used to extracts salience alteration of brain connection of AD in diagnosis as well as in order in order to detect changes in the abnormal convergence of brain which might occur in brain disorders, we generate structure-correlated attributes on graphs.

Our model architecture is comprising of multiple layer graph-CNN networks as cascades of spatial graph convolution →bath normalization →Relu for both encoder-decoder and the discriminator block. We directly trained this model on 96 subjects from preprocess ADNI dataset, including 48 AD and 48 normal control (NC), which describe above and used ADAM [41] optimizer with learning rate set to 10^-4^, momentum of 0.09 during update, mini-batch size of 200 sample that train for 3*10^15^ iterations.

To verify our model, we get two datasets, a real rs-FMRI data from the benchmark ADNI and the other synthetic dataset to develop the illusion correlation matrices to prediction abnormal segmentations.

The synthetic dataset made to further investigation and predict possible construction. This dataset is generated by sampling the Wishart distribution [42] over the average of all ADNI subjects matrices [43] of two AD and NC groups by computing the measure in term of log-likelihood from the classes estimated by the Wishart distribution to show that our model can distinguishes salient nodes or different connection between AD and NC groups, as well as by the synthetic dataset, we detected abnormal area and identified possible state for each defective of brain regions.

Figure 2 depicts the generated graphs by the decoder that fed with given partial graph in the input and several possible solutions on the output. Possible modes of significant correlation between different regions brain from an incomplete graph is produced via synthetic AD dataset. One of the advantages of our approach is to find close convergence between brain regions and predict fitting connections or reconstructing where only partial functional connectivity data is available. Eventually, during a competition to minimize the equation 3, the most appropriate graph is chosen.

**Fig 2.**
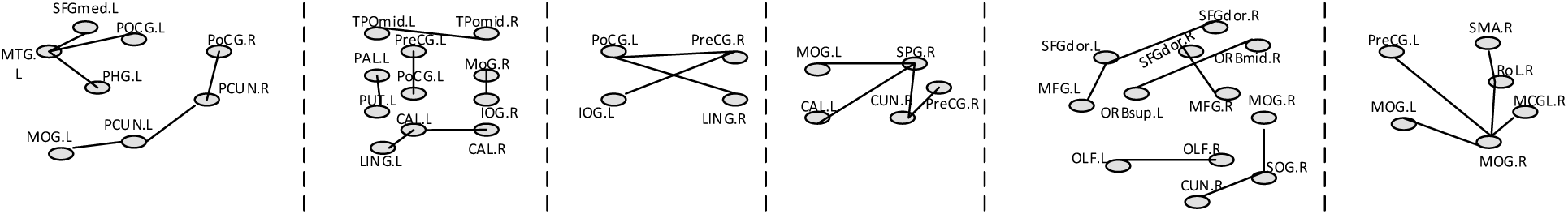
Possible solutions to complete the input partial graph that are revealed by the model. Abbreviation: L,left & R,right;PreCG, Precentral gyrus; SFGmed, Super frontal gyrus, medial;PCUN, Precuneus; CUN, Cuneus; CAL, Calcarine; LING, Lingual gyrus; IOG, Inferior occipital gyrus; MTG, middle temporal gryus; MOG, middle occipital gyrus; ROL, Rolandic opercular; SMA, Supplementary motor area; PoCG, Postcentral gyrus; PHG, Parahippocampal gyrus; SPG, Super parietal gyrus; SFGdor, Dorsolateral of the superior the frontal gyrus; ORBsup, orbital part of the superior frontal gyrus; OLF, Olfactory cortex;SOG, Superior occipital gyrus; TPOmid, Temporal pole, middle part; PAL, lenticular nucleus, pallidum;PUT, lenticular nucleus, putamen; MFG, Middle frontal gyrus, orbital.

To mitigate the excessive cost involved in computing and converging faster, into significate connection of the AD group, we consider larger weights for unusual connection to highlight correlation of these relationships. However, the use of synthetic dataset will cause the results in the *open-world assumption* drive, while it is desirable here for unexpected prediction on the partial connections.

In our future study, based on the analysis above and focus on these topological attributes to extend this work, we will extract important information on the higher-order functional of brain network via semantic functional rich-club organization.

